# A meta-analysis of the stony coral tissue loss disease microbiome finds key bacteria in lesions and unaffected tissue of diseased colonies

**DOI:** 10.1101/2022.09.30.510359

**Authors:** Stephanie M. Rosales, Lindsay K. Huebner, James S. Evans, Amy Apprill, Andrew C. Baker, Anthony J. Bellantuono, Marilyn E. Brandt, Abigail S. Clark, Javier del Campo, Caroline E. Dennison, Naomi E. Huntley, Christina A. Kellogg, Mónica Medina, Julie L. Meyer, Erinn M. Muller, Mauricio Rodriguez-Lanetty, Jennifer L. Salerno, William B. Schill, Erin N. Shilling, Julia Marie Stewart, Joshua D. Voss

## Abstract

Stony coral tissue loss disease (SCTLD) has been causing significant whole colony mortality on reefs in Florida and the Caribbean. The cause of SCTLD remains unknown, with limited concurrence of SCTLD-associated bacteria among studies. We conducted a meta-analysis of SSU 16S ribosomal RNA gene datasets generated by 16 field and laboratory SCTLD studies to find consistent bacteria associated with SCTLD across disease zones (vulnerable, endemic, and epidemic), coral species, coral compartments (mucus, tissue, and skeleton), and disease states (apparently healthy colony tissue [AH], and unaffected [DU] and lesion [DL] tissue from diseased colonies). We also evaluated bacteria in seawater and sediment, which may be sources of SCTLD transmission. Although AH colonies in endemic and epidemic zones harbor bacteria associated with SCTLD lesions, and aquaria and field samples had distinct microbial compositions, there were still clear differences in the microbial composition among AH, DU, and DL in the combined dataset. Alpha diversity between AH and DL was not different; however, DU showed increased alpha diversity compared to AH, indicating that, prior to lesion formation, corals may undergo a disturbance to the microbiome. This disturbance may be driven by Flavobacteriales, which were especially enriched in DU. While Rhodobacterales and Peptostreptococcales-Tissierellales were prominent in structuring microbial interactions in DL. Peptostreptococcales-Tissierellales specifically may contribute to lesion progression through an alpha-toxin. We provide a consensus of SCTLD-associated bacteria both prior to and during lesion progression and identify how these taxa vary across studies, coral species, coral compartments, seawater, and sediment.

## Introduction

Stony coral tissue loss disease (SCTLD) causes focal or multifocal lesions on hard coral colonies (order Scleractinia) leading to exposed skeleton from tissue loss [1, 2].

Affected colony mortality rates can be as high as 99%, but survival is highly dependent on the coral species [3]. While some corals, such as branching Caribbean acroporids, are not impacted by this disease [3], SCTLD nevertheless has a wide host range, affecting over half of Caribbean coral species (~22 species) [2, 3]. This has resulted in a decline in coral species richness, coral cover, and ecosystem function throughout Florida and the Caribbean [3–10].

The cause of SCTLD is currently unknown, but multiple hypotheses of the potential etiology have been proposed, including abiotic stressors [3, 4, 11, 12], viruses [13, 14], bacteria [15, 16], or a combination. SCTLD was first detected in September 2014 off the coast of Miami, Florida, coincident with a dredging project to expand the Miami Port and a coral bleaching event [3, 4]. This led to speculation that heat stress and/or sedimentation may be linked to SCTLD. Evidence to date suggests that thermal stress has either no association with SCTLD [17] or slows disease progression [18], but sediments may contribute to SCTLD transmission [11, 12].

SCTLD is contagious and can be transmitted through the water column [18–20] or through direct coral-coral contact [21], suggesting that it is caused by a biotic source(s) [9, 22]. Viruses have been found in SCTLD-affected corals, although similar virus morphologies and sequences were also detected in apparently healthy corals at similar abundances [13, 14]. Studies have also detected ciliates [1, 22, 23] and endolithic organisms [24] associated with SCTLD, but other eukaryotes have not been associated with this disease.

The most well-studied SCTLD microbial group is the bacterial community, examined using small subunit (SSU)16S rRNA gene analysis [11, 12, 15, 16, 23, 25–29]. It is likely that the bacterial community is important for SCTLD progression, since there is a shift in bacterial composition from healthy corals to diseased corals, and SCTLD lesion progression can be mitigated using antibiotics [22, 30, 31]. Orders such as Rhodobacterales, Rhizobiales, Clostridiales, Alteromonadales, and Vibrionales have been described across many studies, but there have been discrepancies, especially at finer taxonomic levels. Further, a consensus on the key bacteria associated with SCTLD across locations and coral species remains a topic of discussion.

The lack of consensus across studies may be due to biological factors such as coral species-specific microbiomes, the environment in which samples were collected, or other biological variables. However, variability across studies may also arise from different laboratory processing, library preparation, and analytical approaches [32]. In addition, results are often reported at different taxonomic levels such as order [11], family [16], genus [15], and species [25], which can make it difficult to compare across studies. Thus, to better understand SCTLD, a meta-analysis of available SSU 16S rRNA datasets can reduce biases associated with pipelines and reporting strategies. In this study, we examined microbiome datasets from 16 SCTLD studies using a consistent analysis pipeline to determine global patterns and taxa associated with SCTLD.

## Results

### Summary of SCTLD microbiome studies

Initially, datasets were acquired from 17 SCTLD studies, but one study [26] did not pass quality filtering and was removed from the analysis, resulting in 16 SCTLD studies used in this meta-analysis. Additionally, one *Acropora* spp. rapid tissue loss (RTL) disease study was included for comparison of bacteria which may be associated more generally with coral tissue loss diseases (Supplemental Table 1). The combined dataset included 2,425 samples, representing various coral species and environments described below. After the removal of miscellaneous samples such as lab controls, 2,362 samples remained (Supplemental Table 1). Samples from the studies were sequenced using five primer pairs: CS1-515F/CS2-806R [33] with additional 5’ linker sequences [34] (n=79), 515FY [35]/806RB [36] (n=1,219), S-D-Bact-0341-b-S-17/S-D-Bact-0785-a-A-21 [37] (n=31), 515F/806R [33] (n=49), and 515F [33]/Arch806R [38] (n=984; Figure 1A).

**Figure 1.**
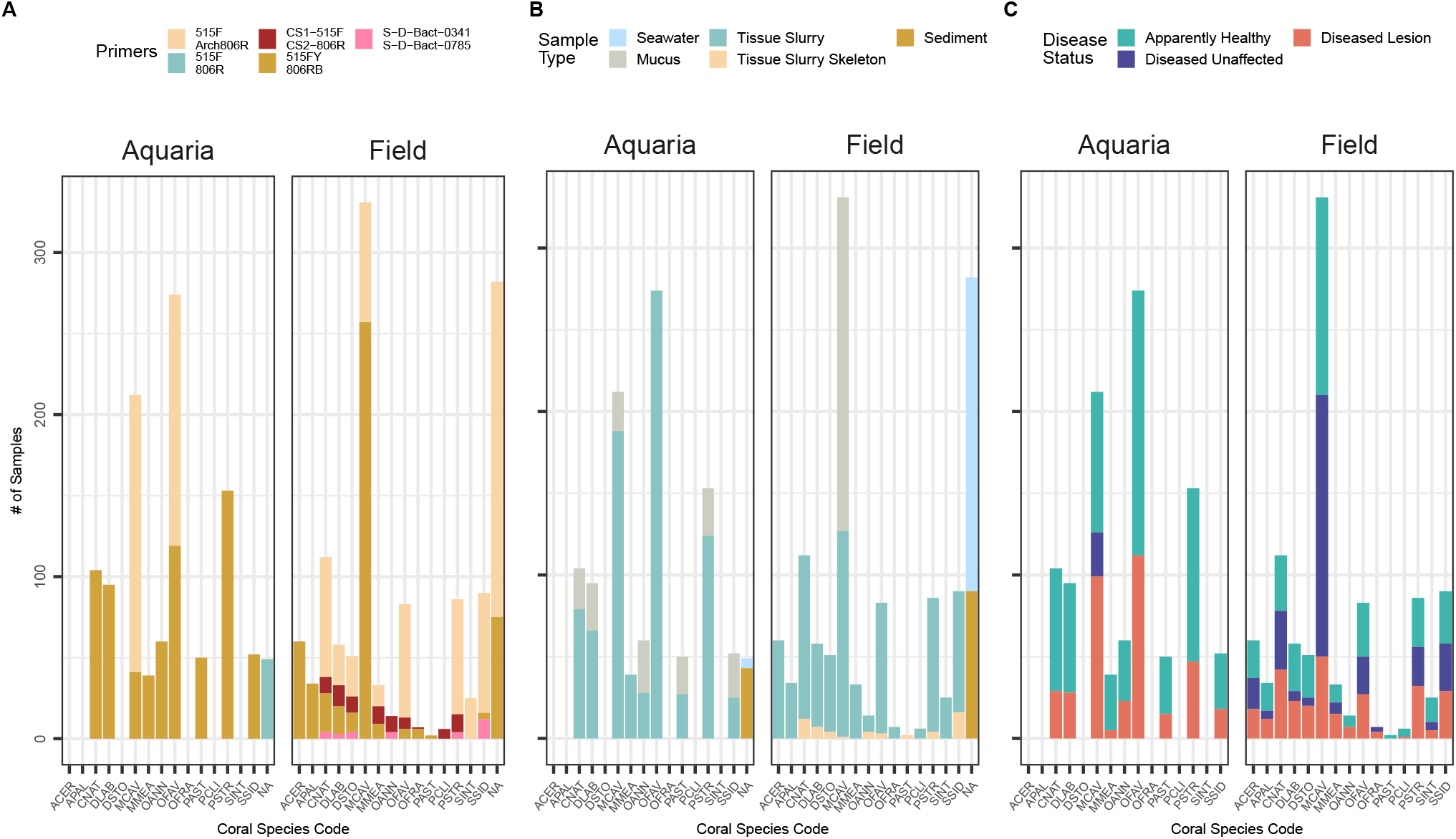
The number of aquaria and field samples for each coral species across (A) SSU rRNA gene primer sets, (B) sample type, and (C) disease state. NAs in (A) and (B) represent sediment and seawater samples. Coral species codes represent the following: *Acropora cervicornis* (ACER), *Acropora palmata* (APAL), *Colpophyllia natans* (CNAT), *Diploria labyrinthiformis* (DLAB), *Dichocoenia stokesii* (DSTO), *Montastraea cavernosa* (MCAV), *Meandrina meandrites* (MMEA), *Orbicella annularis* (OANN), *Orbicella faveolata* (OFAV), *Orbicella franksi* (OFRA), *Porites astreoides* (PAST), *Pseudodiploria clivosa* (PCLI), *Pseudodiploria strigosa* (PSTR), *Stephanocoenia intersepta* (SINT), *Siderastrea siderea* (SSID).

Samples were collected throughout Florida and the U.S. Virgin Islands (USVI). Field samples totaled 1,274, representing 40 sites, and a further 1,088 samples were from aquaria (i.e., laboratory-based experiments; Figure 1). Thirteen SCTLD-susceptible coral species were included, with *Montastraea cavernosa* (MCAV; n=543) and *Orbicella faveolata* (OFAV; n=357) most represented and *Pseudodiploria clivosa* (PCLI; n=6) and *Orbicella franksi* (OFRA; n=7) least represented (Figure 1). Coral samples (n=2,031) were from three compartments: mucus only (n=393), mucus and surface tissue (tissue slurry; n=1,585), and skeleton samples with embedded coral tissue (tissue slurry skeleton; n=53). Seawater (n=198) and sediment (n=133) samples from both the field and aquaria experiments also were included (Figure 1B). Coral samples represented three SCTLD health states: apparently healthy colonies (AH), which was the most represented (n=1,021), followed by lesions on diseased colonies (DL; n=661), and unaffected areas on diseased colonies (DU; n=349; Figure 1C).

### Differences in the microbial composition were found among zones (vulnerable, endemic, and epidemic)

For alpha diversity for AH field-sourced samples, after filtering, 41,504 amplicon sequence variants (ASVs) remained, which were reduced to 15,021 following rarefaction. Among AH samples, there was a slight decrease in Shannon (alpha) diversity from the vulnerable zone (estimated marginal means [emmean]=3.95) to the epidemic zone (emmean=3.70), but this was not significant (Supplemental Fig. 1). For beta diversity, the dispersion was also not different between zones. A PERMANOVA pairwise comparison was significant for all comparisons between zones (p-adjusted [padj] <0.03; Figure 2A), which was adjusted by location given that epidemic samples from the USVI clustered separately from Florida epidemic samples. Differential abundance analysis found 61 ASVs enriched between vulnerable and endemic sites (Figure 2B and Supplemental Table 2). In the endemic zone, the orders Synechococcales (*Cyanobium* PCC-6307; log-fold=12.67) and an uncultured Flavobacteriales (log-fold=9.96) contained ASVs with the highest log-fold change, but the order Flavobacteriales was the group of bacteria with the most enriched ASVs (n=13), followed by SAR11 clade (n=4) and Rhodobacterales (n=3). Fewer ASVs were enriched between the vulnerable and epidemic zones (n=31; Figure 2C and Supplemental Table 3), with the highest log-fold ASV changes found in the orders Burkholderiales (*Delftia;* log-fold=5.84) and Peptostreptococcales-Tissierellales (*Fusibacter;* log-fold=5.65). Like in endemic sites, Flavobacteriales was the group with the most enriched ASVs in the epidemic zone (n=5).

**Figure 2.**
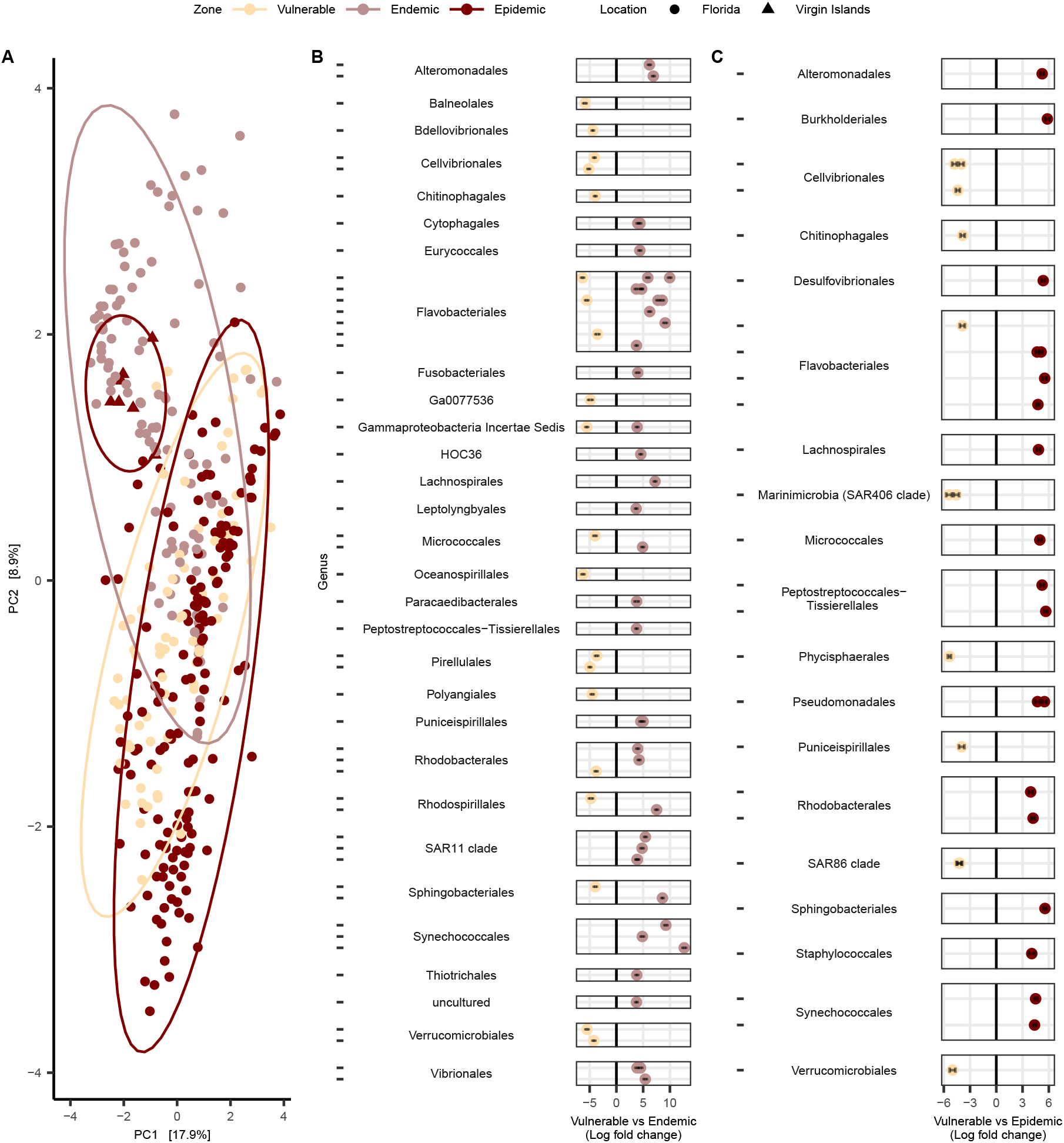
Comparisons among microbial communities of field-sourced apparently healthy (AH) coral colonies across SCTLD zones (vulnerable, endemic, and epidemic) in Florida and the U.S. Virgin Islands in (A) beta diversity (centered log-ratio transformed and plotted with a Euclidean distance), and differential abundance analysis in (B) vulnerable vs endemic zones, and (C) vulnerable vs epidemic zones. ASVs are grouped by genus (represented by dashes) on the y-axis and then by order, and only ASVs with a padj<0.001, a W statistic >90, and a log-fold change <-2 and >2 were visualized. AH samples from the three coral compartments (mucus, tissue slurry, and tissue slurry skeleton) were included and *Acopora* spp. samples were excluded from the analysis.

### Disease state showed the lowest correlation to beta diversity

Microbial dispersion was found to be different across primers, study, biome, year, all coral species, and sample type (Permutest: *p*<0.01; Figure 3). A PERMANOVA test for differences between microbial composition was also significant across all factors, with coral species having the highest correlation (R^2^=0.21; Figure 3E) and disease state showing the lowest correlation (R^2^=0.04).

**Figure 3.**
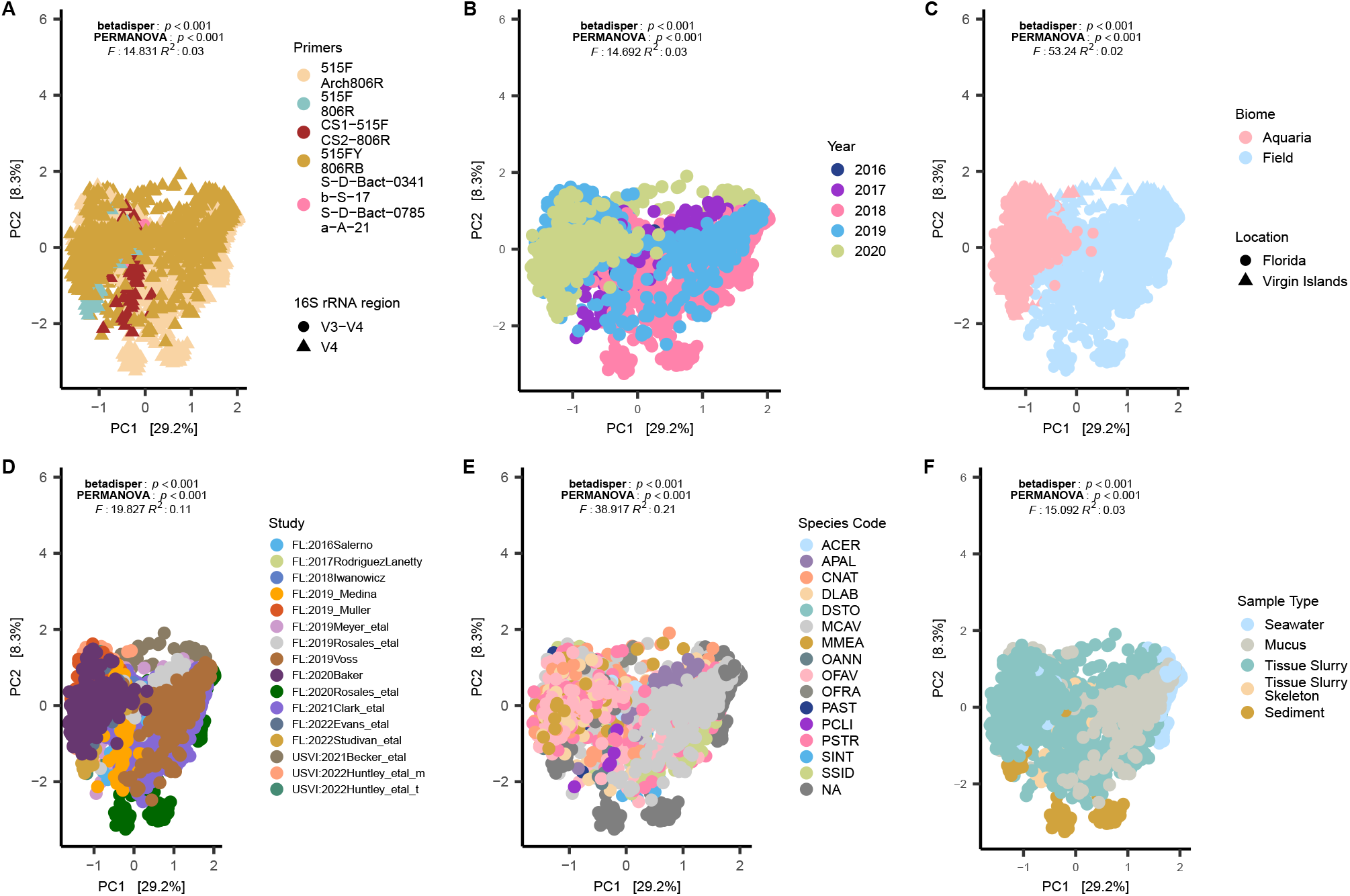
Microbial beta diversity of all coral species (SCTLD-susceptible corals and *Acropora* spp.) and sample types (coral, sediment, and seawater) show differences within and between microbial communities in (A) SSU 16S rRNA gene primers, (B) year, (C) biome, (D) study, (E) coral species, and (F) sample type. All plots were centered log-ratio transformed and visualized with a Euclidean distance. The NAs in (E) represent sediment and seawater samples; coral species codes are defined in Figure 1 legend.

### Differences found between aquaria and field samples based on disease state

Biome (i.e., aquaria and field) had the largest correlation to principal component 1 (PC1, R^2^=0.73; Supplemental Fig. 2) than other tested metadata factors, and showed a distinct separation when visualized (Figure 3C). This was also evident even in sediment and seawater samples that were collected in aquaria studies, which clustered with coral samples from aquaria studies and not with field sediment and seawater samples. Given this pattern, SCTLD-affected corals (with the removal of *Acropora* spp.) were first analyzed combined (i.e., both aquaria and field) and then subsequent analysis was divided by biome to identify potential differences between the two. When both biomes were combined (Figure 4A), AH microbial communities were the most highly dispersed compared to both DU (padj<0.01) and DL (padj<0.02), but DU vs DL were not different. Pairwise PERMANOVA was significant for all comparisons (padj<0.03 each): AH vs DU (R^2^=0.07), AH vs DL (R^2^=0.03), and DU vs DL (R^2^=0.13; Figure 4A). Among aquaria samples (Figure 4B), the dispersion was lower in DU vs both DL and AH (padj<0.0001 each) but was similar in AH vs DL. Like the combined samples, all aquaria samples were different in the pairwise PERMANOVA (R^2^=0.03, padj<0.03 each; Figure 4B). In field samples (Figure 4C), dispersion was lowest in DL compared to both AH and DU (padj<0.0001 each) but there was no difference between DU vs DL. All pairwise PERMANOVA comparisons were significant in the field samples: AH vs DU (R^2^=0.01, padj<0.03), AH vs DL (R^2^=0.03, padj<0.01), and DU vs DL (R^2^=0.08, padj<0.03; Figure 4C).

**Figure 4.**
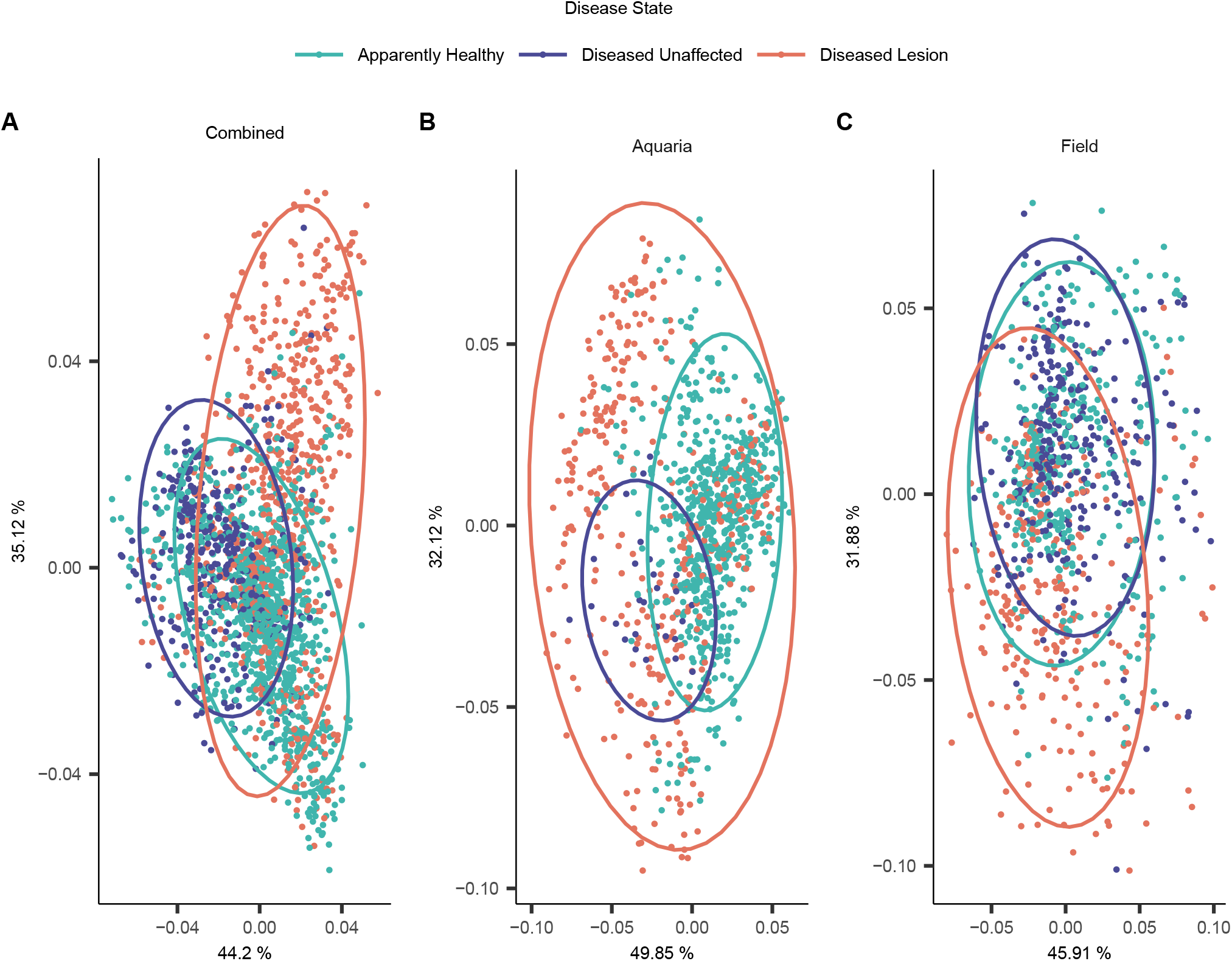
Microbial differences in coral disease state among apparently healthy colonies (AH), and unaffected (DU) and lesion (DL) areas on diseased colonies in beta diversity of (A) both aquaria and field samples (“Combined”), (B) aquaria, and (C) field samples only. Samples from *Acropora* spp. were excluded and the three coral compartments (mucus, tissue slurry, and tissue slurry skeleton) were included in this analysis.

Samples were also evaluated for alpha diversity by disease state in each biome. After quality filtering and rarefaction across disease states, 39,513 ASVs remained. For aquaria and field samples combined, pairwise comparisons showed differences in Shannon diversity for AH vs DU and DL vs DU (padj<0.0001 each) but not AH vs DL, with mean alpha diversity lowest in DL (emmean=3.42) and highest in DU (emmean=3.85; Supplemental Fig. 3A). In aquaria samples only, there were no differences in Shannon diversity by disease state, likely due to the low sample size of DU (n=27, Supplemental Fig. 3B). In field samples, only DU vs DL was different (padj<0.01) with DU also showing the highest mean (emmean=3.90) and DL the lowest (emmean=3.63; Supplemental Fig. 3C) alpha diversity.

When comparing differences in mean relative microbial abundances within disease states across biomes, AH samples differed between aquaria and field (Supplemental Fig. 3D): the orders Rhodobacterales (14.20±5.2%) and Cytophagales (9.02±12.32%) were dominant in aquaria samples, but in field samples, the dominant orders were Flavobacteriales (5.75±2.15%) and Synechococcales (3.77±5.88%). Like AH aquaria samples, DU aquaria samples had the highest mean relative abundances in Rhodobacterales, but at a much lower percentage (1.06±3.81%). The DU field samples were also similar to their AH counterparts, showing the highest relative abundances in Flavobacteriales (6.43±1.89%) and Synechococcales (4.45±6.26%). In the DL samples, both aquaria and field samples were dominated by Rhodobacterales, but the aquaria samples had a higher relative abundance of Rhodobacterales (15.34±6.84%) than samples from the field (6.61±4.12%). As with aquaria AH samples, Cytophagales (3.28±11.22%) were also the second most relatively abundant order in DL aquaria samples but were not dominant in field DL samples. Peptostreptococcales-Tissierellales was a dominant DL member at similar mean relative abundances in both aquaria (3.21±6.40%) and field samples (3.79±9.06%; Supplemental Fig. 3D).

### Indicator taxa were detected across coral compartments, seawater, and sediment

The combined three coral compartments (mucus, tissue slurry, and tissue slurry skeleton), from both field and aquaria, yielded a total of 109 differentially abundant ASVs between AH vs DU (Figure 5A and Supplemental Table 4). DU mucus samples showed the highest log fold change compared to AH in the orders Flavobacteriales (NS5 marine group; log-fold=6.33) and Synechococcales (*Cyanobium* PCC-6307; log-fold=6.19), with Flavobacteriales having the most enriched ASVs (n=9). Similarly, DU tissue slurry samples were most enriched in Synechococcales (*Synechococcus* CC9902; log-fold=20.04) and Flavobacteriales (NS5 marine group; log-fold=12.71), with Flavobacteriales having the most enriched ASVs (n=9). Tissue slurry skeleton sample comparisons of AH vs DU identified no ASVs enriched in DU. In addition to coral compartment samples, ASVs enriched in AH and DU samples were also present within sediment and seawater samples (Figure 5B).

**Figure 5.**
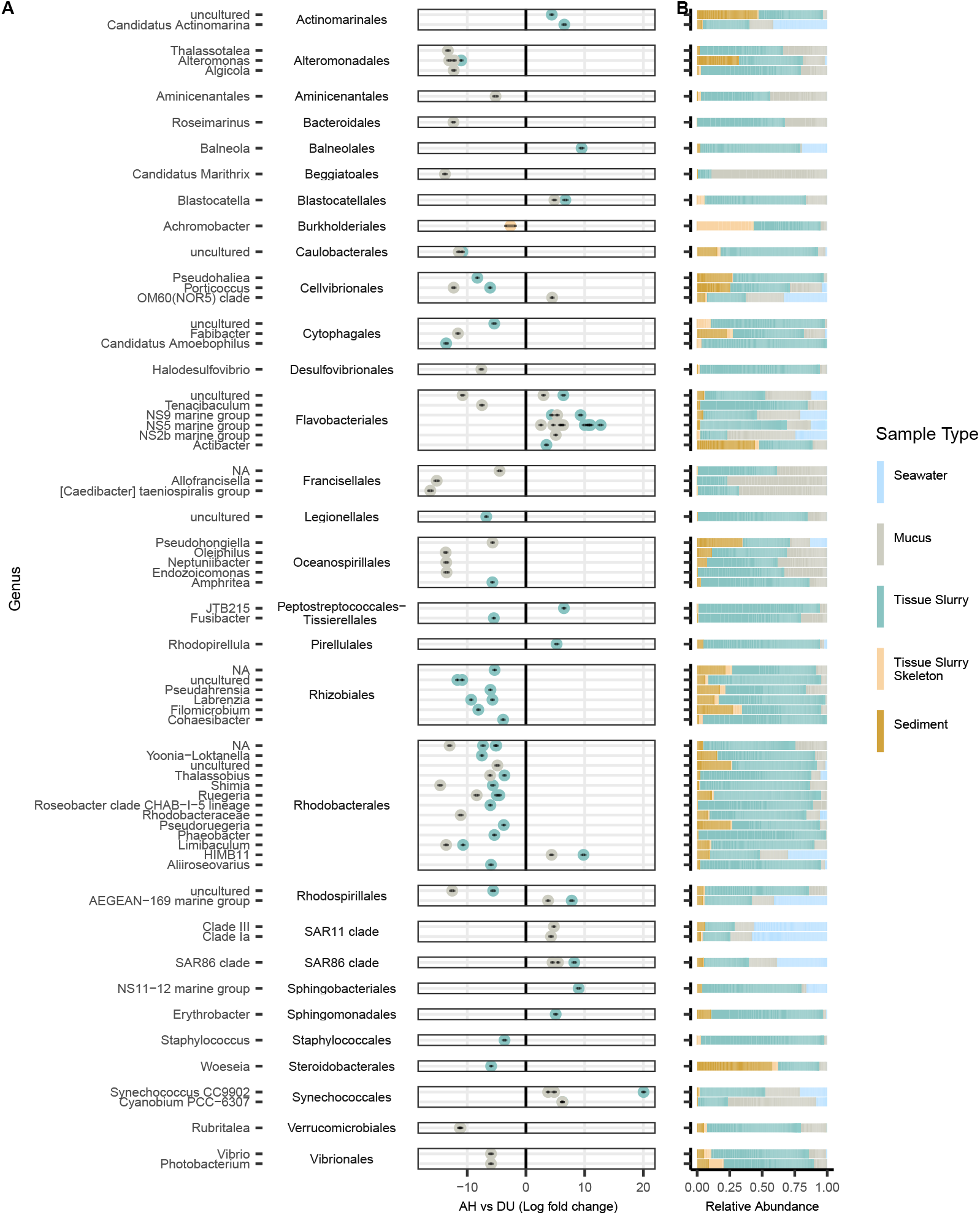
Microbial ASVs associated with unaffected areas on diseased colonies (DU). Differential abundances between (A) apparently healthy (AH) vs DU. The y-axis depicts ASVs grouped by genus and then by order. Only ASVs with a padj<0.001, W statistic >90, and a log-fold change <-1.5 and >1.5 were visualized. Coral compartments (i.e., mucus, tissue slurry, and tissue slurry skeleton) were included and *Acropora* spp. were excluded from this analysis. (B) The relative abundance of taxa enriched in AH and DU by sample type.

When OFAV and MCAV (the most-sampled coral species) were removed from the analysis, similar patterns were still identified in beta diversity and differential abundance when compared to the analysis of all SCTLD-susceptible species (Supplemental Fig. 4A and B). For DU, 18 (35.3%) ASVs were shared between the two analyses (i.e., with vs without OFAV and MCAV), but more unique ASVs were found enriched in the analysis without OFAV and MCAV compared to the analysis that included all SCTLD-susceptible corals. Still, the two analyses shared more enriched bacterial families compared to the number that was enriched only within each individual analysis.

The three combined coral compartments yielded fewer differentially abundant ASVs in AH vs DL (n=79; Figure 6A and Supplemental Table 5) compared to AH vs DU (Figure 5A). In DL mucus samples, ASVs from the orders Desulfovibrionales (*Halodesulfovibrio;*log-fold=13.96) and Rhodobacterales (*Shimia;* log-fold=13.18) were the most enriched, and Rhodobacterales had the most enriched ASVs overall (n=8). In DL tissue slurries, the ASVs with the highest enrichment were two Rhodobacterales from an uncharacterized genus (log-fold=15.77) and one from the genus *Tropicibacter* (log-fold=13.46). Rhodobacterales was also the order with the most enriched ASVs in DL compared with AH tissue slurries (n=14), followed by Peptostreptococcales-Tissierellales (n=6). Among tissue slurry skeleton samples, only one ASV was enriched in DL (Burkholderiales, *Achromobacter;* log-fold=1.49). ASVs enriched in DL were also found in sediment and seawater (Figure 6B). In DL, the differential abundance analysis without OFAV and MCAV compared to that with all SCTLD-susceptible coral species showed that the majority of enriched ASVs were shared (n=25; 39.1%) between the two analyses (Supplemental Fig. 4C).

**Figure 6.**
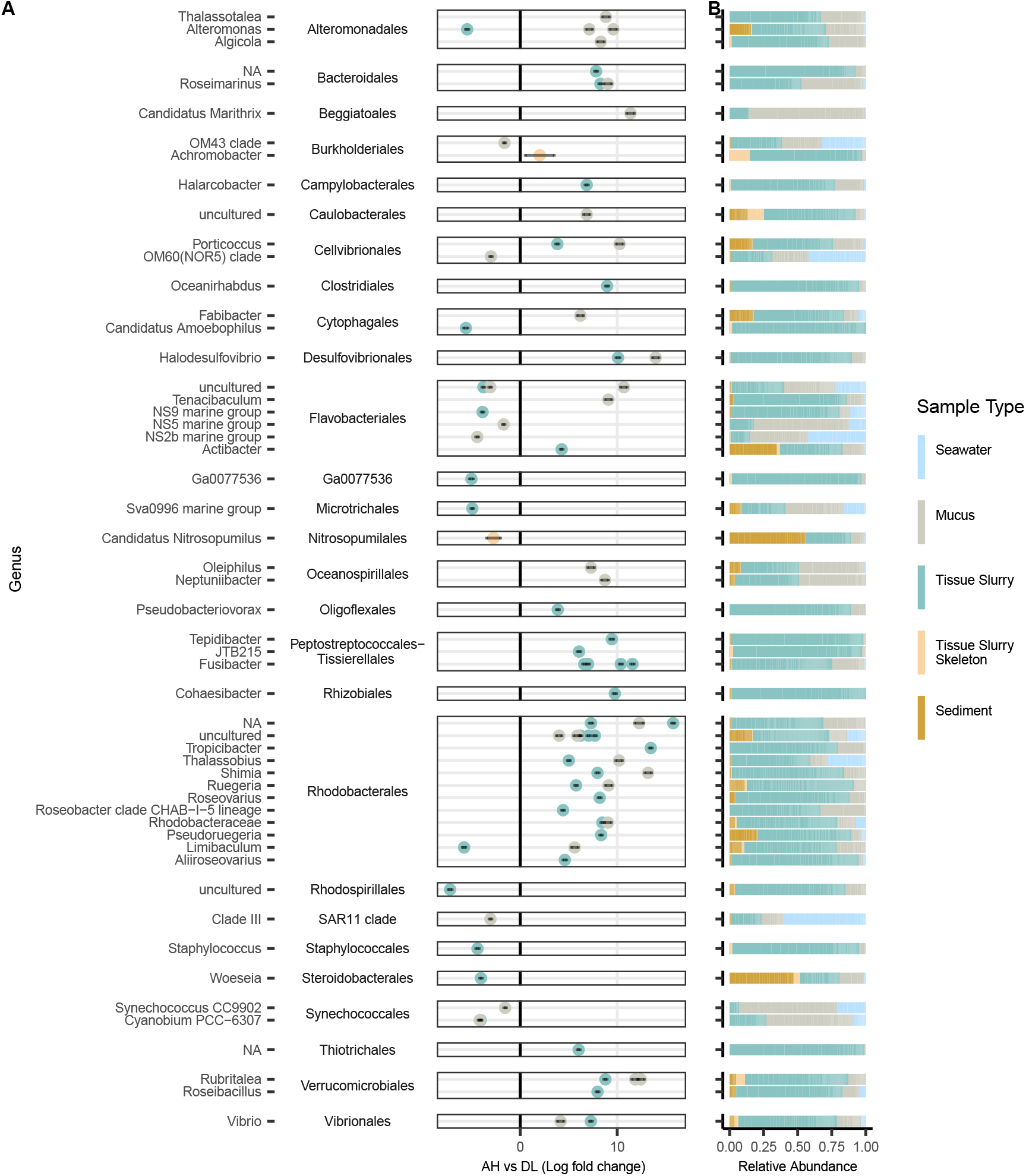
Microbial ASVs associated with lesions on diseased colonies (DL). Differential abundances between (A) apparently healthy (AH) vs DL. The y-axis depicts ASVs grouped by genus and then by order. Only ASVs with a padj<0.001, W statistic >90, and a log-fold change <-1.5 and >1.5 were visualized. Coral compartments (i.e., mucus, tissue slurry, and tissue slurry skeleton) were included and *Acropora* spp. were excluded from this analysis. (B) The relative abundance of taxa enriched in AH and DL by sample type.

### The presence of indicator taxa varied across coral species and studies

Six coral species were represented by a high number of samples (n>76 samples each), and all ASVs only enriched in DU were found within all of those species. The seven coral species with lower sampling frequencies (n<76 each) varied in the numbers of DU-enriched ASVs present (Supplemental Fig. 5A). For example, *Dichocoenia stokesii* (DSTO) contained all DU-enriched taxa, and *Stephanocoenia intersepta* (SINT) had all genera present but one, which belonged to Flavobacteriales. In comparison, *Pseudodiploria clivosa* (PCLI) had the fewest DU-enriched taxa (n=3) among the coral species. Four orders were not present in *Acropora* spp. samples and included: Blastocatellales, Pirellulales, Sphingobacteriales, and Peptostreptococcales-Tissierellales. Across studies, the order Sphingobacteriales was not found in any aquaria study but was found in 50% of field studies (Supplemental Fig. 5B). Additionally, no aquaria study had representatives from all DU-enriched taxa, but four field studies were found to have all taxa. The two studies with the fewest representatives were studies that used V3-V4 primers (Supplemental Table 1).

The ASVs enriched only in DL were also present in all high-frequency coral species, while none of the low-frequency coral species had all of the DL-enriched taxa (Supplemental Fig. 6A). PCLI possessed the fewest DL-enriched genera (n=9) followed by *Orbicella franksi* (OFRA; n=15). More DL-enriched orders (n=11) were absent from *Acropora* spp. Corals (n=4) than DU-enriched orders; the DL orders not present in *Acropora* were: Bacteroidales, Beggiatoales, Burkholderiales, Cellvibrionales, Clostridiales, Desulfovibrionales, Oligoflexales, Peptostreptococcales-Tissierellales, Rhizobiales, Thiotrichales, and Verrucomicrobiales. Across studies, three had all the DL-enriched orders (all aquaria studies), and the fewest orders were present in those which used V3-V4 primers (Supplemental Fig. 6B), as with the DU-enriched orders.

### Alphaproteobacteria and Clostridia were found as important players in SCTLD bacterial community interactions

In a network analysis of co-associated ASVs, a total of nine modules were identified, with two that were significantly and positively correlated to AH (R^2^=0.26 and 0.1), three to DU (R^2^=0.12, 0.31, and 0.46), and four to DL (R^2^=0.17, 0.22, 0.46, and 0.47; Supplemental Fig. 7). The modules with the highest positive correlation to each disease state had 134 (AH; blue), 158 (DU; green), and 146 (DL; pink) co-abundant ASVs (Supplemental Fig. 7) and were used for undirected network analysis (Figure 7).

**Figure 7.**
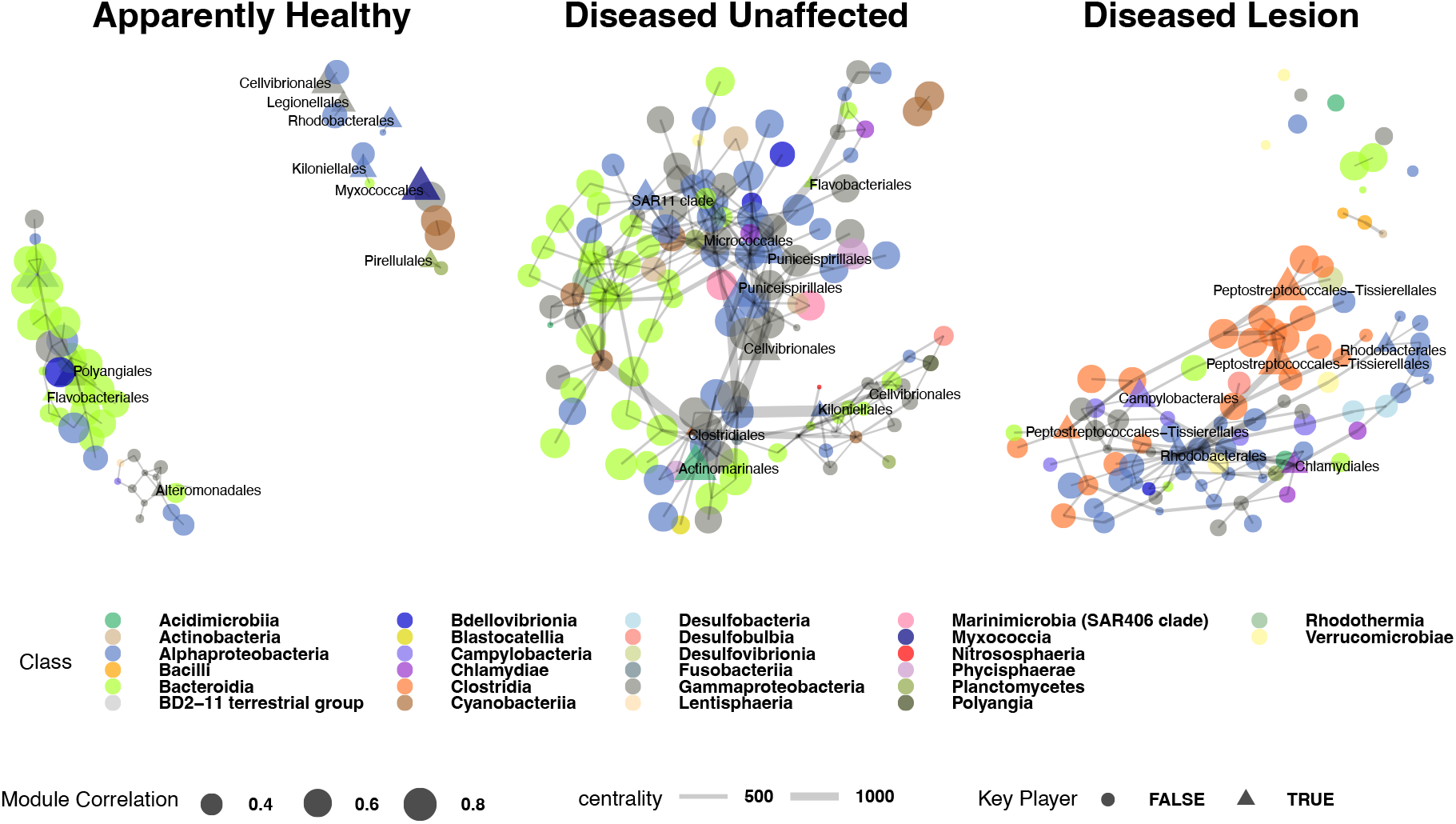
Co-occurrence networks of bacteria from weighted correlation network analysis (WGCNA) modules (Supplemental Fig 7) among apparently healthy colonies (AH), and unaffected areas (DU) and lesions (DL) on diseased colonies. The nodes represent amplicon sequence variants (ASV), which are sized by the ASV’s correlation value to its respective module. A triangle and label of the bacteria order denote that a node is a “key player.” The width of the edges corresponds to centrality, with thicker edges representing higher centrality. Samples from the three coral compartments (i.e., mucus, tissue slurry, and tissue slurry skeleton) were included in the analysis, and *Acropora* spp. samples were excluded from this analysis.

Although AH had the second largest module, the network was smaller than both DU and DL, with only 56 ASV nodes and 59 edges (connections between nodes). DU had the largest network, with 138 nodes and 293 edges, followed by DL, with 123 nodes and 204 edges. In AH, the node with the most neighbors (n=7) was from the class Polyangia (order Polyangiales), which was also considered in the AH network to be a key player (i.e., provides cohesiveness, connectedness, and is embedded in a network; Figure 7).

The two nodes with the highest correlation to the blue weighted correlation network analysis (WGCNA) module (Supplemental Fig. 7) were from the class Bacteroidia (Chitinophagales; R^2^=0.88 and 0.87).

In the DU network, highly connected nodes included three Alphaproteobacteria (SAR11 clade [n=16], Rhodobacterales [n=11], and Rhodospirillales [n=11]; Figure 7). Alphaproteobacteria were among the classes assigned as key players, but additional key players included: Cyanobacteria, Bacteroidia, and Polyangia. The nodes most highly correlated to their respective WGCNA modules were SAR86 clade (R^2^=0.88) and Rhodospirillales (R^2^=0.88).

The DL network had nodes with the most neighbors compared to AH and DU and was driven by Alphaproteobacteria (two Rhodobacterales nodes [n=22 and n=16], and Rhizobiales [n=9]), and Bacteroidia (Flavobacteriales [n=12]; Figure 7). While Alphaproteobacteria (Rhodobacterales and Rhizobiales) were found as key players in DL, Flavobacteriales were not. Additional key players in DL included Clostridia, Chlamydiae, and Campylobacteria. The class Clostridia had the highest correlations to the DL pink WGCNA module (Peptostreptococcales-Tissierellales; R^2^=0.77 and Lachnospirales R^2^=0.76; Supplemental Fig. 7). The most prevalent classes in DL networks were Alphaproteobacteria (n=39; mainly Rhodobacterales, n=29) and Clostridia (n=23; mainly Peptostreptococcales-Tissierellales, n=13).

### The top microbial functional pathways were more enriched in DL compared to AH and DU

To identify differences in the potential microbial function between disease states, we used the SSU16S rRNA gene for functional predictions. There was a total of 6,307 differently abundant (padj<0.05) Kyoto Encyclopedia of Genes and Genomes (KEGG) pathways identified across AH (n=2,482), DU (n=1,403), and DL (n=2,422). Of the top 10 KEGG pathways, three were enriched in DU and six in DL (Supplemental Fig. 8A). The most enriched pathway in DU was 4-hydroxybutyrate dehydrogenase (effect size=0.25), and in DL was phospholipase C/alpha-toxin (effect size=0.97). A total of 392 differentially abundant MetaCyc pathways were found across AH (n=148), DU (n=104), and DL (n=139). Out of the top 10 pathways, nine were enriched in DL and one in DU (Supplemental Fig. 8B). Biotin biosynthesis II was the most enriched pathway in DL (effect size=0.80), and ADP-L-glycero-β-D-manno-heptose biosynthesis in DU (effect size=0.05).

## Discussion

We used a crowdsourcing approach of both unpublished and published data to better understand stony coral tissue loss disease (SCTLD) across zones of disease spread (vulnerable, endemic, and epidemic), coral species, biomes (field vs aquaria), and studies to provide a more informed consensus on SCTLD-associated bacteria. We identified potential changes to coral microbiomes based on the length of time the disease had been present in the area (i.e., epidemic vs endemic zones). We also found differences in alpha and beta diversity by coral disease state: apparently healthy colonies (AH), and unaffected areas (DU) and lesions (DL) on diseased colonies. Furthermore, DU and DL showed unique sets of enriched bacteria, with DL microbiomes particularly structured by Rhodobacterales and Peptostreptococcales-Tissierellales interactions.

### Apparently healthy field-sourced coral microbiome composition differed among SCTLD zones

To understand if SCTLD alters the microbiome of visibly healthy corals on SCTLD-affected reefs, we examined AH corals within three disease zones: vulnerable, epidemic, and endemic. Although there were no differences in alpha diversity and dispersion among zones, microbial beta diversity and enriched microbial taxa were different among zones, as previously documented [11]. AH corals in the endemic and epidemic zones harbored SCTLD-associated microbes such as Alteromonadales, Vibrionales, Peptostreptococcales-Tissierellales, and Rhodobacterales, potentially indicating they were actively combating or showing signs of resistance to the disease. Flavobacteriales was the group with the most enriched taxa in endemic and epidemic AH corals, which is notable because Flavobacteriales were also detected in both DU and DL and are known to associate with corals under stressful conditions [39]. As AH corals showed no outward signs of lesions, members of Flavobacteriales may represent initial members of the SCTLD microbiome. However, a better understanding of the specific species or strains of Flavobacteriales present in both healthy and diseased corals may explain their enrichment in different health states as Flavobacteriales was also found at high abundances in AH.

### SCTLD aquaria studies may change microbial dynamics compared to field studies

We found that both biomes (aquaria and field) had distinct microbial compositions, which has been reported previously [40]. Despite this, we detected a microbial composition shift in disease states in both biomes, but field samples had a stronger beta diversity correlation in AH vs DL comparisons. Furthermore, there were notable differences in the relative abundances of certain taxa between biomes. For example, while Rhodobacterales were dominant members in both biomes, they were found at higher relative abundances in aquaria compared to field samples. Because Rhodobacterales are primary surface colonizers in marine waters, including surfaces such as glass [41] aquarium environments may provide conditions that particularly enrich Rhodobacterales over other bacterial taxa. Additionally, aquaria showed high relative abundances of Cytophagales in AH and DL, but this taxon was not a top abundant order in field samples. An aquaria coral challenge study with *Vibrio coralliilyticus* also showed an enrichment of both Rhodobacterales and Cytophagales [42], further indicating that aquarium conditions may select for these two taxa. Interestingly, Peptostreptococcales-Tissierellales, a bacterial group that appears to be important within SCTLD lesions, was present at similar relative abundances in DL in both biomes and thus may be less susceptible to laboratory artifacts than Rhodobacterales.

Notably, there were two aquaria experiments designed to limit the ‘microbial background noise’ from the field by using sterilized seawater [29] or sterilized sediment [12] and then incubating the chosen medium with healthy or diseased corals. The resulting bacterial communities from the incubated seawater and sediment clustered with aquaria coral samples rather than with field seawater and sediment samples, and show that these inoculums likely take on the host microbial community. While field sediment and seawater samples showed distinct community separation from field coral samples, these samples still showed enrichment of some ASVs found in field DU and DL tissues, indicating potential transfer of microbes between diseased corals and their environment, which may result in continued transmission of SCTLD [11].

### Unaffected tissues on diseased colonies were enriched with Flavobacteriales and Synechococcales

This meta-analysis provides a comprehensive list of important microbial taxa in SCTLD across three coral compartments (mucus, tissue slurry, and tissue slurry skeleton) and three disease states (AH, DU, DL). DU areas on coral colonies are of interest as they may represent an initial disturbance from SCTLD to the microbial community with potentially fewer secondary and saprophytic bacteria. It is unknown if SCTLD is a localized or systemic condition, but histological studies have found internal SCTLD lesions in DU tissue prior to lesion formation on the colony surface [21, 24]. We found that DU samples had the highest alpha diversity and a distinct microbial composition, further suggesting that SCTLD also causes disruptions in the microbiome prior to surface lesion formation. Compared to AH, DU becomes particularly enriched with Flavobacteriales (class Bacteroidia) and Synechococcales (class Cyanophyceae), and these taxa formed strong connections to the class Alphaproteobacteria from the orders SAR11 and Rhodobacterales. However, SAR11 interactions may only be important in the mucus, which was the only compartment in which they were enriched.

Of note, the increase in DU alpha diversity could be partly driven by the lack of standardization of DU samples. One study collected DU samples from colonies in which DL tissues were treated with antibiotics, potentially disrupting the DU microbiome [43]. The remaining studies consistently sampled DL at the lesion margin, but the DU samples varied in distance collected from the lesion. This may be driving the diversity detected, as the DU microbial community is known to change with distance from the lesion, with samples closest to the lesion possessing more SCTLD-associated taxa than those farther away [15]. A standardized definition of DU should be employed to maximize the utility of these samples. Regardless, the majority of DU-enriched taxa are likely not primary pathogens, as they were also found in the *Acropora* spp. rapid tissue loss (RTL) study. However, one Blastocatellales and Peptostreptococcales-Tissierellales ASV each were not found in the RTL study but were prevalent in 73% of the SCTLD studies, and therefore could be specific to SCTLD. Although these two ASVs have a 100% sequence similarity to bacteria found within black band disease (accession MH341639; [44]), and a paling necrosis study (GU200211.1; [43]), respectively, the studies took place outside of the Caribbean and thus the ASVs could belong to bacteria newly introduced to the area.

### Rhodobacterales and Peptostreptococcales-Tissierellales were key structural components of microbial interactions in disease lesions

There was no clear transition from AH to DU to DL in alpha diversity, and AH and DL alpha diversity values were similar. It may be difficult to capture a general microbial alpha diversity response to SCTLD across coral species, as alpha diversity values are highly species-specific [28]. However, there were differences in microbial composition between AH and DL. In DL, the microbial community transitioned into an enrichment of Rhodobacterales and Peptostreptococcales-Tissierellales, which belong to the classes Alphaproteobacteria and Clostridia, respectively. Clostridia are anaerobic [46] and while Rhodobacterales are generally aerobic [47] they can thrive in anoxic conditions [48]. This suggests that as the disease state transitions from DU to DL, the lesion may progress to anoxic conditions, as seen in black band disease [49]. However, presently it cannot be determined if that is a result of actions by the bacteria or if their enrichment is based on the shifting lesion environment [50]. Nonetheless, these two classes showed the highest connectivity and presence in the network analysis.

Across SCTLD microbiome studies, Rhodobacterales has been reported as highly abundant in all except one [26], and while Peptostreptococcales-Tissierellales has been found enriched in some studies [15, 16, 27, 28], Clostridia has been significantly enriched in all of them [11, 15, 16, 23, 26–28]. Rhodobacterales may be more generally associated with coral diseases, as many of the taxa associated with SCTLD were also found in the *Acropora* spp. RTL study. In contrast, Peptostreptococcales-Tissierellales was not found in RTL and a BLAST search of these ASVs showed that only one had 100% similarity to a sequence in the database, from a study that examined soil polluted by crude oil [49]. The rest were less than 94.31% similar to the NCBI 16S rRNA database, suggesting these taxa may be unique to SCTLD. Analysis of inferred functional traits showed that Clostridia taxa may have important roles in lesion progression through pathways such as phospholipase C / alpha-toxin, a toxin found in Clostridia such as *Clostridium perfringens* [52] and a top pathway predicted in DL in this study. Phospholipase C / alpha-toxin is a metalloenzyme, dependent on zinc ions, which through lipid signaling degrades eukaryotic cell membranes, potentially resulting in necrosis [52]. Thus, Peptostreptococcales-Tissierellales could be contributing to tissue loss in SCTLD via an alpha-toxin. Overall, the high network connectivity and inferred functional potential of toxin production suggest that Clostridia may have a particularly important role in SCTLD bacterial interactions and lesion progression. Therefore, promising future directions for SCTLD microbiome research could include developing enrichment media for Clostridia [53] and then conducting knockout gene studies of the alpha toxin genes [54].

### Future SCTLD studies may consider sampling less-studied coral species

In this meta-analysis, only half of the coral species impacted by this disease were evaluated [2]. While we found consistent bacterial enrichment between analyses with and without the two most frequently sampled coral species, coral species was found to be the main factor driving microbial community structure. Therefore, analyzing representatives of all susceptible coral species could be especially important in further narrowing down the microbial taxa specific to SCTLD. Future studies could consider including coral species with no or low sampling representation in their permits to enable opportunistic sampling, which when pooled together in a collaborative analysis such as this, may yield meaningful results.

## Conclusions

This is the largest microbiome meta-analysis ever conducted on a single coral disease. We found differences in the microbiomes of apparently healthy (AH) corals between SCTLD zones (vulnerable, endemic, and epidemic). In endemic and epidemic zones, AH corals may have acquired SCTLD-associated bacteria, potentially representing a compromised health state or resistance. We also identified that dominant taxa varied depending on whether the samples were collected away from the lesion (DU) or near the lesion (DL) on a colony with SCTLD. In DU samples, Flavobacteriales and Synechococcales were the dominant taxa, but in DL Rhodobacterales and Peptostreptococcales-Tissierellales were dominant and were key taxa in structuring microbial networks. This indicates that there is a shift of dominant bacterial taxa during disease progression and implies the lesion tissue may become anoxic. Specifically, during lesion progression, Peptostreptococcales-Tissierellales may be involved in tissue loss by lysing coral and symbiont cells through the phospholipase C / alpha-toxin pathway. Peptostreptococcales-Tissierellales taxa also appear to be more specifically associated with SCTLD and not coral disease generalists, as some of the ASVs found here have not been reported in other coral diseases.

Our findings convey the need for further focus on the transition of bacterial taxa from DU to DL and characterization of the role of Peptostreptococcales-Tissierellales in lesion progression. A key aspect of this future work could be the inclusion of a wider assortment of coral species and compartments to better clarify the mechanisms of SCTLD. In addition, more holistic studies are needed to understand SCTLD. The bacterial community appears to play a role in SCTLD, but other members of the holobiont (i.e., viruses and Symbiodiniaceae) may be contributing to this change. Combining multiple methods such as culturing, metagenomics, metatranscriptomics, and microscopy could help better clarify the microbial disease dynamics in SCTLD.

## Methods

### Obtaining studies

To acquire small subunit (SSU) 16S rRNA datasets for this meta-analysis, an email was sent on July 14, 2020, and July 23, 2020, to the hosts of the coral-list listserv and the SCTLD Disease Advisory Committee (DAC) email list, requesting scientists to share unpublished SCTLD-associated microbiome datasets. To allow for comparisons of microbiomes between a past Caribbean coral disease to the novel SCTLD outbreak, a rapid tissue loss (RTL) disease study in *Acropora palmata* (APAL) and *Acropora cervicornis* (ACER) comprising apparently healthy (AH) samples, inoculated AH samples, and inoculated diseased samples [55], also was included in some analyses. This particular study was selected because *Acropora* spp. reportedly are not susceptible to SCTLD and the study used V4 primers [3]. In total 17 studies were analyzed, 16 from SCTLD and one from an *Acropora* spp. RTL study (Supplemental Table 1).

Study authors were requested to complete a pre-formatted metadata file to facilitate comparisons of data across studies. Requested metadata included sample handling information (e.g. source laboratory, sample collector) and ecological information (e.g. source reef name, coordinates, zone, water temperature, and coral colony measurements). SCTLD zones included vulnerable (where the disease had not yet arrived), endemic (where the initial outbreak of the disease had moved through and no or few active lesions were observed on colonies), and epidemic (where the outbreak was active and prevalent across colonies of multiple species). Invasion zone sites, where the disease was newly-arrived but not yet prevalent, were grouped within the epidemic zone for consistency across studies and simplicity of analysis. Some metadata required standardization of units and not all metadata were available across all studies.

However, in all field-collected samples, all sampling dates and reef site information were available, enabling the completion of SCTLD disease zone metadata for Florida studies by referencing the Coral Reef Evaluation and Monitoring Project, Disturbance Response Monitoring, and SCTLD boundary reconnaissance databases. For USVI, zones were assigned based on the USVI Department of Planning and Natural Resources SCTLD database (https://dpnr.vi.gov/czm/sctld/).

### Bioinformatics to process sequence data

Each sequencing run was imported to QIIME2-2022.2 [56, 57] and processed individually. The datasets were divided into two distinct pipelines: (1) data that targeted the 16S rRNA gene V4 region of Bacteria and/or Archaea and (2) data that targeted the V3-V4 region of Bacteria and/or Archaea. For V4 datasets, the data were processed with cut-adapt to remove sequencing primers corresponding to the respective study [58]. In total, three 515F primers that targeted the V4 region of the 16S rRNA gene were used across studies (‘5-GTGCCAGCMGCCGCGGTAA-3’ [n=1,033] [33], ‘5-GTGYCAGCMGCCGCGGTAA-3’ [n=1,219] [35], and ‘5-ACACTGACGACATGGTTCTACAGTGCCAGCMGCCGCGGTAA-3’, [n=79] [33, 34]; Supplemental Table 1). Next, the data were processed with DADA2 for quality control and denoising using a max error rate of three [59]. Although all runs were paired-end reads, the V4 samples were processed as single-end reads and the forward reads were truncated at 130 base pairs (bp) with the DADA2 program. The error rates, truncation, and single-end options were selected based on the quality and sequence length (Supplemental Table 1) of the lowest quality reads across all datasets. The two V3-V4 datasets (n=31 samples) were processed with the cut-adapt program, which was used to select forward sequences that contained sequences similar to the 515F primers used in the V4 studies. The forward primer 515FY [35] was used as the target sequence using a 0.4 error rate to allow for some differences in bases. The selected sequences were then processed with DADA2 and truncated at 240 bps with a max error rate of one. After, if studies had multiple Illumina sequencer runs, they were first merged together, and then all studies were merged into one count table and sequence file. The vsearch cluster-features-de-novo function was then used to cluster the data by 99% similarity [60]. The classify-consensus-vsearch option was then used for taxonomy assignments with the SILVA-138-99 database [61]. The data were then filtered to remove mitochondria and chloroplast reads.

### Alpha diversity

Shannon diversity metrics were generated with the phyloseq function rarefy_even_depth with option replace=TRUE, and a minimum sequence depth for a sample of 1000. Prior to rarefaction, taxa with a sum of zero across the subsetted data were removed. Two sets of alpha diversity analyses were run: (1) evaluated differences across the three zones (vulnerable, endemic, and epidemic) in field-sourcws apparently healthy (AH) corals, and (2) evaluated differences across disease states (AH, unaffected tissue [DU], and lesion tissue [DL] on a diseased colony) in SCTLD-susceptible corals (i.e., without *Acropora* spp.). Significance was tested with linear mixed models with the R packages lme4 v1.1.21 [62], and emmeans v1.4.3.1 [63], and for pairwise comparisons Tukey’s HSD was used. For zones and disease states, coral species was used as a random effect.

### Beta diversity

The data were imported into R v4.0.5 and converted into a phyloseq object [64]. ASVs were removed if they were present less than four times in 20% of the samples. The filtered count table was transformed using centered log-ratio (CLR) with the package microbiome [65]. Beta diversity was analyzed with the package VEGAN 2.5.4 [66] and the filtered CLR-transformed table. The function vegdist was used to generate dissimilarity indices with a Euclidean distance. To identify significant differences among groups, a Permutational Multivariate Analysis of Variance (PERMANOVA) was used with the function adonis2 with 999 permutations, using a Euclidean distance. The function betadisper was used to calculate group dispersion, which was then tested for significance with the function Permutest.

Differences in beta diversity for field samples were evaluated in apparently healthy (AH) corals across three zones (vulnerable, endemic, and epidemic), and blocked by location (Florida vs USVI). In addition, pairwise group comparison was assessed from betadisper output using the Tukey’s HSD function. The PERMANOVA output was also tested for pairwise comparisons with the function pairwise.adonis and adjusted with a Bonferroni correction [67]. Furthermore, all samples (including *Acropora* spp., sediment, and seawater) were also evaluated for beta diversity differences in primers, year of collection, biome (field and aquaria), studies, coral species, and sample type (seawater, mucus, tissue slurry, tissue slurry and skeleton, and sediment). These factors were also correlated to principal components (PCs) using the R package PCAtools 2.5.15, and the functions pca and eigencorplot were used to remove the lowest 10% of the variance and to correlate the data and test for significance, respectively.

SCTLD-susceptible coral samples (i.e., without *Acropora* spp., sediment, and seawater) were also evaluated for beta-diversity. Both biomes (aquaria or field) were examined together and also separately. The matrices were generated with QIIME2-2021.11 with the plugin DEICODE, which runs a robust Aitchison Distance – a method that is not influenced by zeros in the data [68]. This was then evaluated for dispersion, differences in microbial composition between groups, and pairwise comparisons. DEICODE was also applied to the data without the two most prevalent corals species, *Orbicella faveolata* (OFAV) and *Montastraea cavernosa* (MCAV), to see if the same pattern was evident in disease states with and without these coral species.

### Differential abundance analysis

The program Analysis of Compositions of Microbiomes with Bias Correction (ANCOM_BC) was used to identify differentially abundant microbial taxa [69]. ANCOM_BC was used with the global test option and the results were considered significant if the false discovery rate adjusted p-value (padj) was <0.001 and if the W statistic was >90. Field-sourced AH samples were tested for differential abundance among zones (vulnerable, endemic, and epidemic), and SCTLD-susceptible coral samples (without *Acropora* spp.) were evaluated for differences in disease state (AH, DU, and DL). For SCTLD-susceptible corals, the data were parsed by the three coral compartments (mucus, tissue slurry, and tissue slurry skeleton). Sistinct ANCOM_BC analyses were run for each compartment due to the relatively low sample size of tissue slurry skeleton samples compared to the two other compartment types. The taxa were further evaluated if they had a log-fold change between −1.5< and >1.5. The ASVs that were significantly enriched were used to identify the relative abundance of the ASVs across sample types. In addition, those enriched only in either DU or DL were used to identify the presence or absence of each ASV in coral species and study per biome. The same ANCOM_BC analysis was repeated without MCAV and OFAV to evaluate if the two dominant coral species in our meta-analysis were driving the enriched bacteria.

### Network analysis

To identify ASVs that co-associate in AH, DU, and DL samples, CLR-transformed counts were used for weighted correlation network analysis (WGCNA) with the WGCNA 1.70-3 R package [70]. The network was constructed unsigned with the following parameters: power=3, minimum module size=12, deep split=2, and merged cut height=0.25. The eigenvalues were correlated to AH, DU, and DL using Pearson correlation with the R function cor. The highest correlation in each disease state was then selected for network construction using the R package SpiecEasi 1.0.5 [71]. The network was then constructed as previously reported [11]. Briefly, the Stability Approach to Regularization Selection (StARS) [72] model was chosen along with the method Meinshausen-Bühlmann’s neighborhood selection [73]. For StARS, 100 subsamples were used with a variability threshold of 10^-3^. The centrality (node importance) was evaluated [74] using the functions centrality_degree (neighbors=the number of adjacent edges or neighbors) and centrality_edge_betweenness (centrality=the number of shortest paths going through an edge) [75]. The package influenceR 0.1.0. [76] selected important ASVs in each network and assigned the top “key players” [77], which were labeled with their respective orders.

### Functional prediction profiles

To infer the functional potential of 16S rRNA gene data among AH, DU, and DL, the program Phylogenetic Investigation of Communities by Reconstruction of Unobserved States (PICRUSt2) was used in QIIME2-2021.11 [78]. Only SCTLD-susceptible corals were evaluated and only ASVs that were present in at least 100 samples were selected. The picrust2 full-pipeline was used with the hidden state set to “mp” and the placement tool to place sequences into a tree set to “epa-ng.” The outputs were predicted metagenomes for Kyoto Encyclopedia of Genes and Genomes (KEGG [79]) orthologs and MetaCyc pathway [80] abundances. To assess the differential abundance of these outputs among disease states, the R package Maaslin2 was utilized [81]. For both KEGG and MetaCyc tests, data were log-transformed, a random effect was set to coral species, and the data were subsequently analyzed with a linear model. In the KEGG assessment, the minimum abundance=0.05 and the minimum prevalence=0.1. There were no minimums set for the MetaCyc test due to the lower number of pathways found in MetaCyc. The top 10 predicted pathways were selected based on values with the lowest padj and effect sizes <-0.5 and >0.5. These pathways were manually annotated on KEGG and MetaCyc websites.

## Supporting information

Supplemental Fig. Line 8

## Acknowledgments

Any use of trade, firm, or product names is for descriptive purposes only and does not imply endorsement by the U.S. Government. We would like to thank the Disease Advisory Committee Pathogen ID/Microbiome sub-team which spawned the “Sweet 16S” team that led to this paper. SMR was supported by National Oceanic and Atmospheric Administration (NOAA) Office of Oceanic and Atmospheric Research (OAR) omics initiative. LKH was supported by Environmental Protection Agency South Florida Initiative grants X7-00D66417-0 and X7-01D00820-0. JSE, CAK, and WBS were supported by the USGS Ecosystems Biological Threats & Invasive Species Research Program. AA was supported by National Science Foundation awards 1938112 and 2109622 and a NOAA OAR Cooperative Institutes award. EMM was supported by a Mote Eminent Scholarship. ENS and JDV were supported by Florida Department of Environmental Protection, Professional Association of Diving Instructors, and the Harbor Branch Oceanographic Institute Foundation.

## Data availability statement

Sequence data available on NCBI are listed in Supplemental Table 1. All other datasets are available upon request. The unfiltered ASV counts table, taxonomy table, ASV sequences, and code to conduct this analysis are publicly available at https://github.com/srosales712/SCTLD_microbiome_meta_analysis.

