## Supplementary figures and images for "A meta-analysis of the stony coral tissue loss disease microbiome finds key bacteria in lesions and unaffected tissue of diseased colonies"

### Supplemental Fig. 1

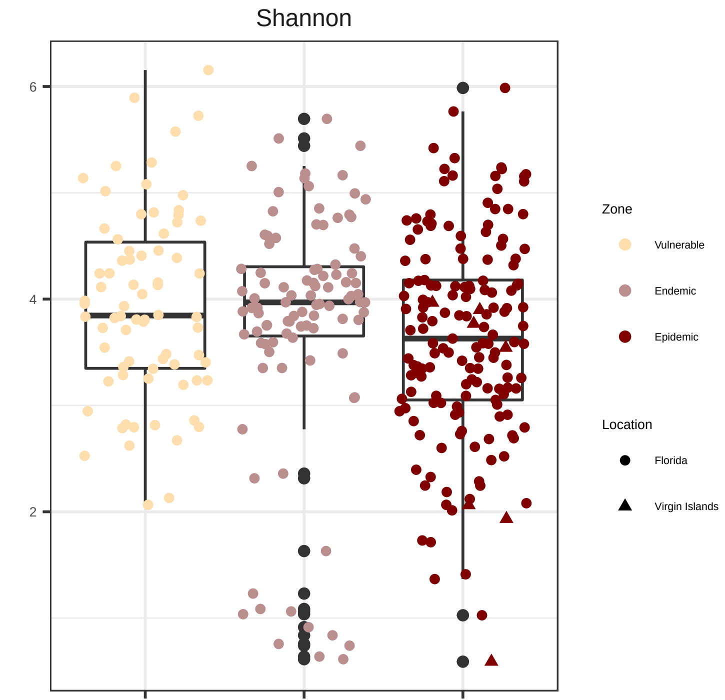

### Supplemental Fig. 2

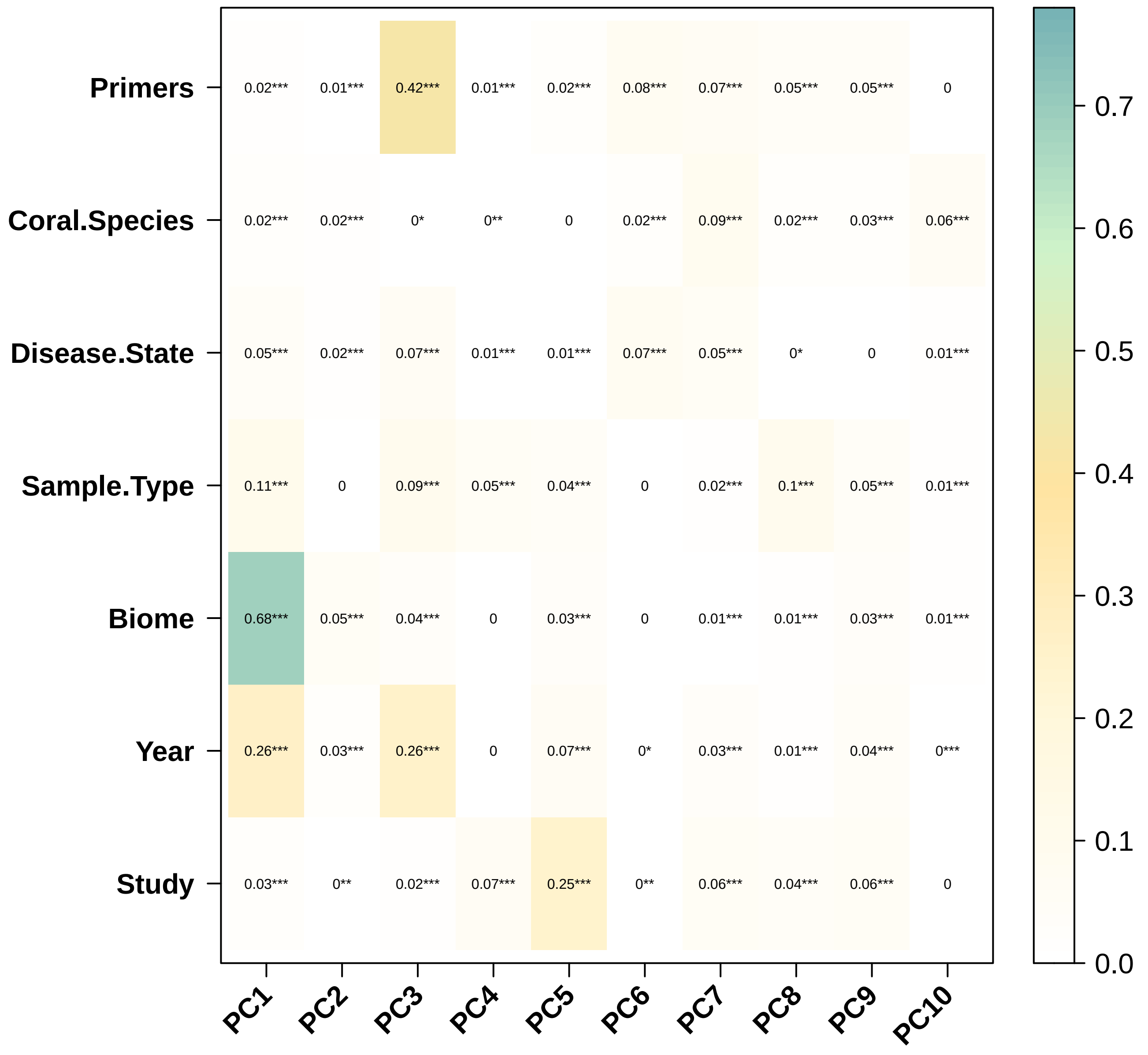

### Supplemental Fig. 3

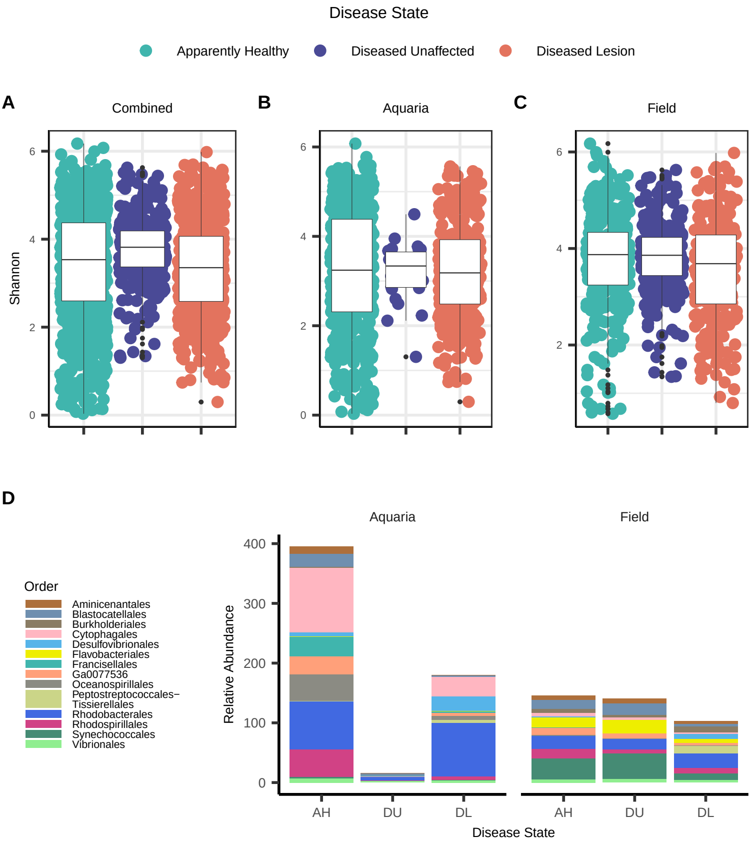

### Supplemental Fig. 4

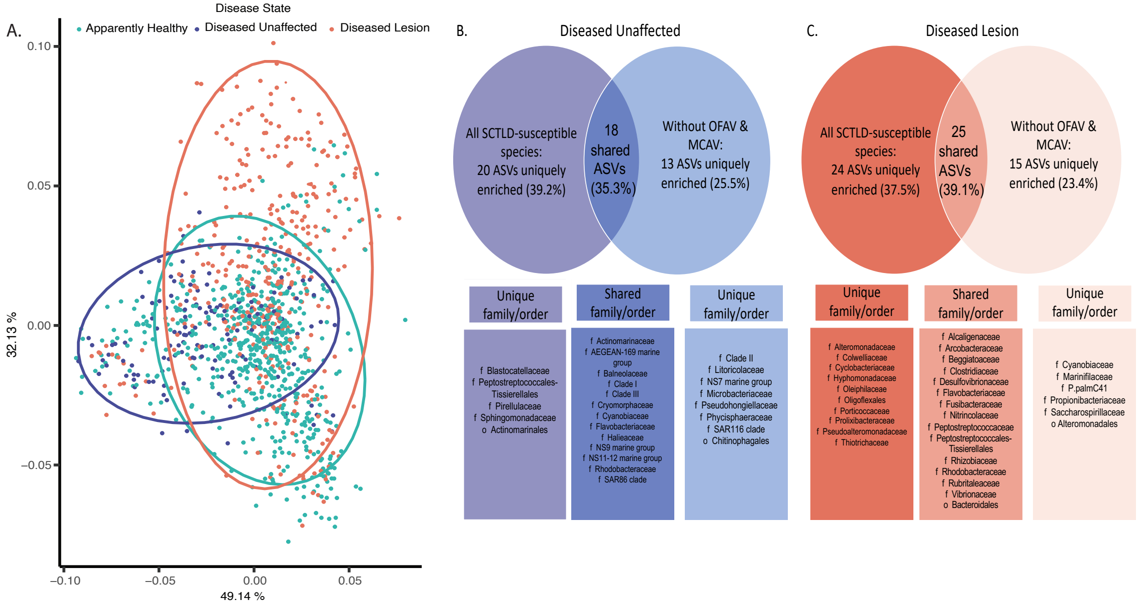

### Supplemental Fig. 5

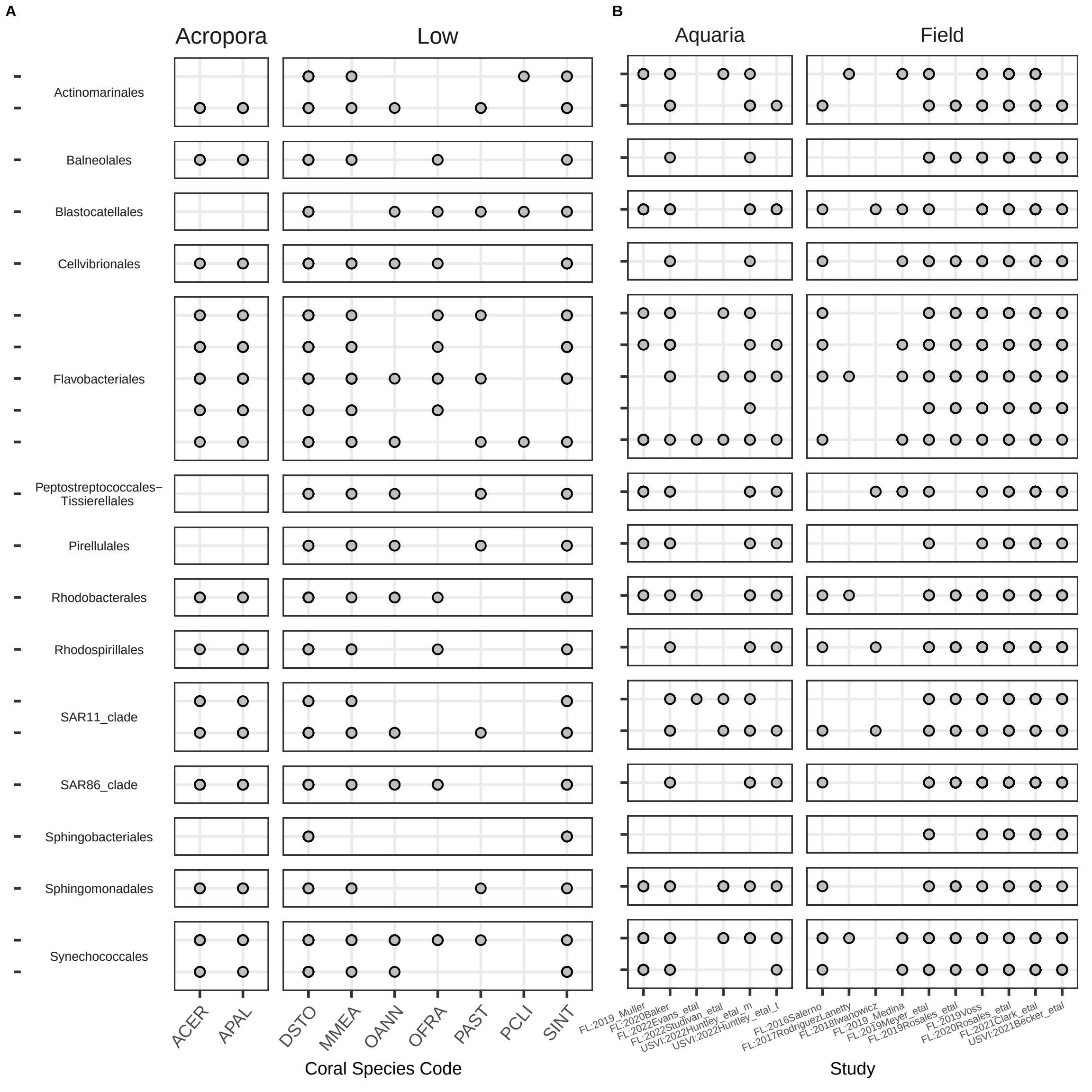

### Supplemental Fig. 6

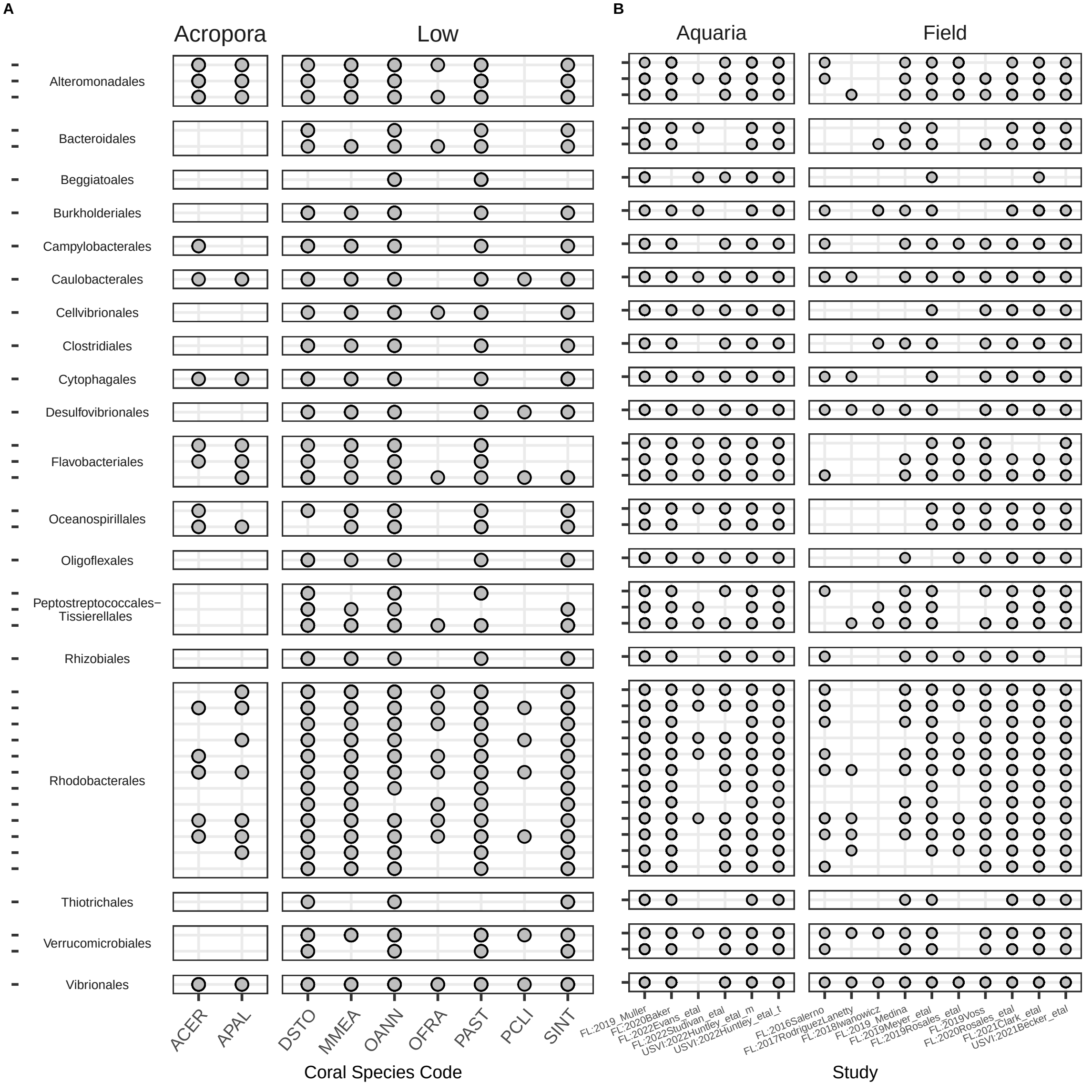

### Supplemental Fig. 7

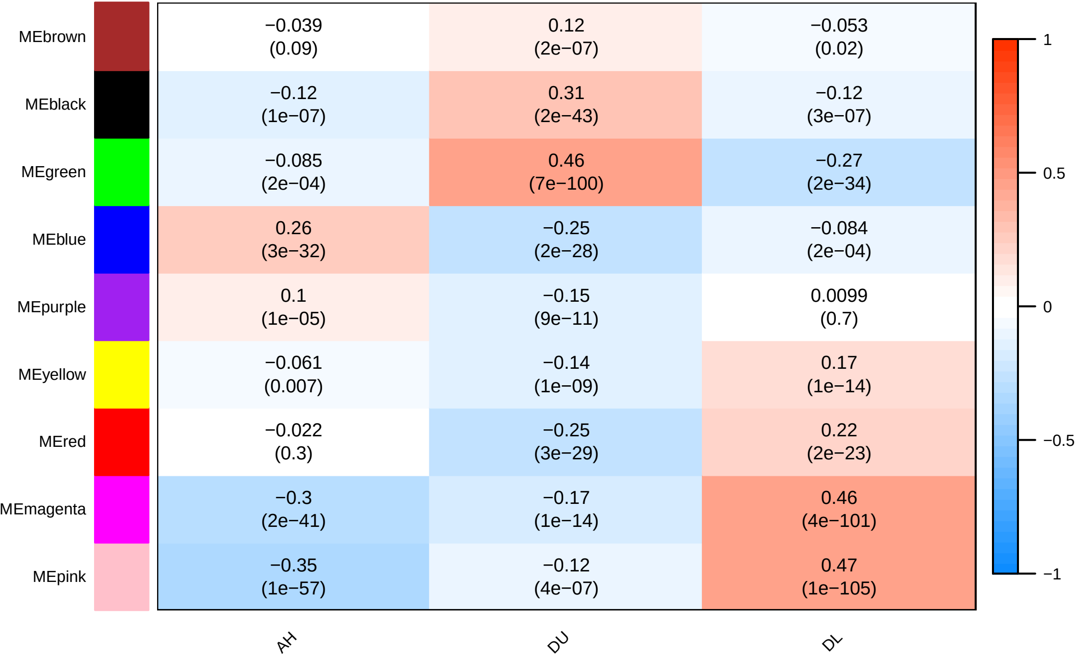

### Supplemental Fig. Line 8

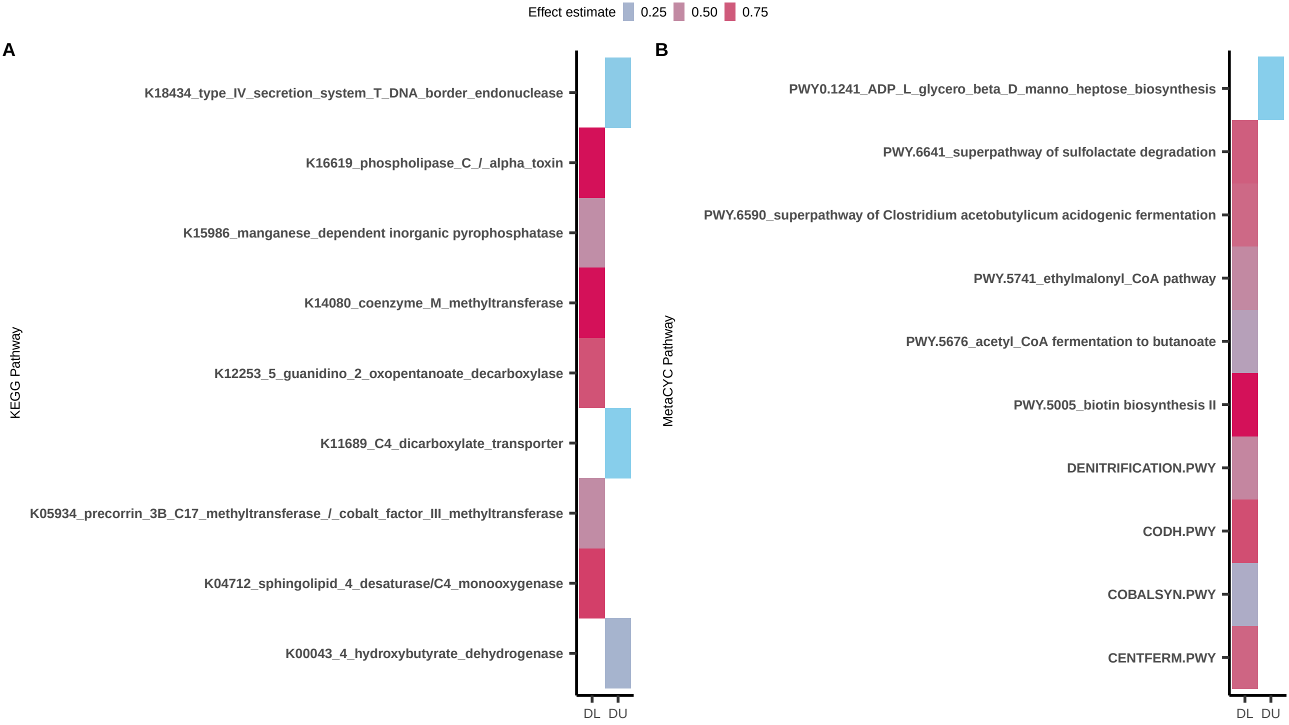
